# A Web-based software toolkit for accessible and best-practice machine learning analyses in biomedical research

**DOI:** 10.64898/2026.06.05.730487

**Authors:** Paulo Lyra, Junhao Qiu, Khai Dang, Alyssa Pybus, Isis Narvaez-Bandera, Maansi Singh, Qiang Gu, Luke Sargent, Allison Creason, Jeremy Goecks

**Author notes:** Equal Contributions (Paulo Lyra, Junhao Qiu).

## Abstract

Machine learning is increasingly central to biomedical research, but using machine learning well often requires substantial computational expertise and methodological care to produce high-quality results. To make machine learning tools more accessible to biomedical researchers while supporting best-practice approaches, we developed the Galaxy Learning and Modeling (GLEAM) software toolkit. GLEAM enables researchers to perform supervised machine learning analyses through a set of web-based, code-free software tools for tabular, image, and multimodal biomedical datasets. GLEAM standardizes data partitioning, model selection, training, evaluation, and reporting, helping researchers apply machine learning with greater rigor and consistency. GLEAM runs on the Galaxy computational workbench and uses Galaxy’s core features to make all analyses accessible, reproducible, and scalable. We validated GLEAM on three biomedical tasks: predicting patient response to immunotherapy, skin lesion classification, and cancer recurrence prediction. Across these tasks, GLEAM produced highly accurate predictive models and improved transparency, reproducibility, and rigor.

---

Machine learning (ML) has become essential to modern biomedical data research. Several of the most cited papers in this century describe ML methods^1^, and review articles across many biomedical fields highlight both methodological advances and practical applications of ML^2–4^. ML is used for tasks ranging from automated image analysis^5^ and imputing missing values^6^ to modeling multimodal omics of cellular function^7^ and clinical applications such as disease classification, risk stratification, and outcome prediction^8,9^. Despite this broad impact, ML can be difficult to use well^10^. Effective use requires methodological knowledge, including which ML models are applicable for different data types and how to evaluate them well^11^. It also requires computational knowledge to use software libraries and computing resources in ways that produce robust, complete, well-documented analyses^12^.

Modern software libraries and analysis platforms have been developed to provide high-level components for doing ML. General-purpose frameworks, such as Scikit-Learn^13^, PyTorch^14^, and TensorFlow^15^, provide ready-to-use ML models and data preprocessing. Meanwhile, automatic ML (AutoML) frameworks, including PyCaret^16^ and AutoGluon^17^, can automate model selection, training, and evaluation. However, these libraries are largely focused on empowering software developers and computational researchers rather than the biomedical research community. Thus, scientists still must define splits, assemble pipelines, manage compute resources, and document settings and results across disconnected components^18,19^. This imposes a high degree of complexity for using ML, especially for researchers without informatics expertise. As a result, ML-focused biomedical analyses remain difficult to perform and vulnerable to avoidable errors in data preprocessing, model selection, and evaluation^20^. These challenges reduce the impact of ML and the validity of ML-based results in biomedical research.

To address these challenges and gaps, we developed the Galaxy Learning and Modeling (GLEAM) software toolkit. GLEAM provides comprehensive code-free and best-practice supervised machine learning for tabular, image, and multimodal biomedical data. Using only a web browser and GLEAM, scientists can perform high-quality and scalable machine learning from start to finish.GLEAM uses intelligent defaults and best-practice ML libraries to simplify machine learning for scientists while promoting robust modeling and evaluation. GLEAM produces comprehensive evaluation reports that summarize model performance and features driving model predictions. GLEAM is built on the popular Galaxy^21–23^ computational workbench, which is used by thousands of scientists worldwide. Galaxy integration powers GLEAM by providing an accessible user interface for its ML tools, complete reproducibility of all modeling, access to computing resources such as GPUs, and integration with thousands of other software tools. To demonstrate GLEAM’s capabilities, we use it to analyze public datasets for three different biomedical applications. We show that GLEAM provides highly accurate performance and illustrate how consistent inputs, explicit split handling, and report-guided evaluation support thorough and best-practice use of ML in biomedical research. A website summarizing GLEAM, with links to its tools, tutorials, and source code, is available at https://goeckslab.github.io/gleam/.

## RESULTS

### GLEAM provides a web-based user interface for rigorous and scalable machine learning

The GLEAM software platform implements five principles for rigorous biomedical machine learning (ML): accessibility, integration, reproducibility, scalability, and best practices (**Fig. 1a**). GLEAM includes three machine learning software tools that are named to reflect the type of data each learner uses for input and modeling: `Tabular Learner`, `Image Learner`, and `Multimodal Learner`. Each learner supports classification and regression and uses an established machine learning software library selected for the target input data modality (**Fig. 1b; Supplementary Table 1 and 2**). All GLEAM learners are accessible via a user-friendly web-based interface, are completely reproducible, and can scale to large analyses. To promote comprehensive evaluation of ML results, learners produce an interactive report that includes 15+ performance metrics and interpretability plots and tables **(Supplementary Table 3)**.

**Figure 1.**
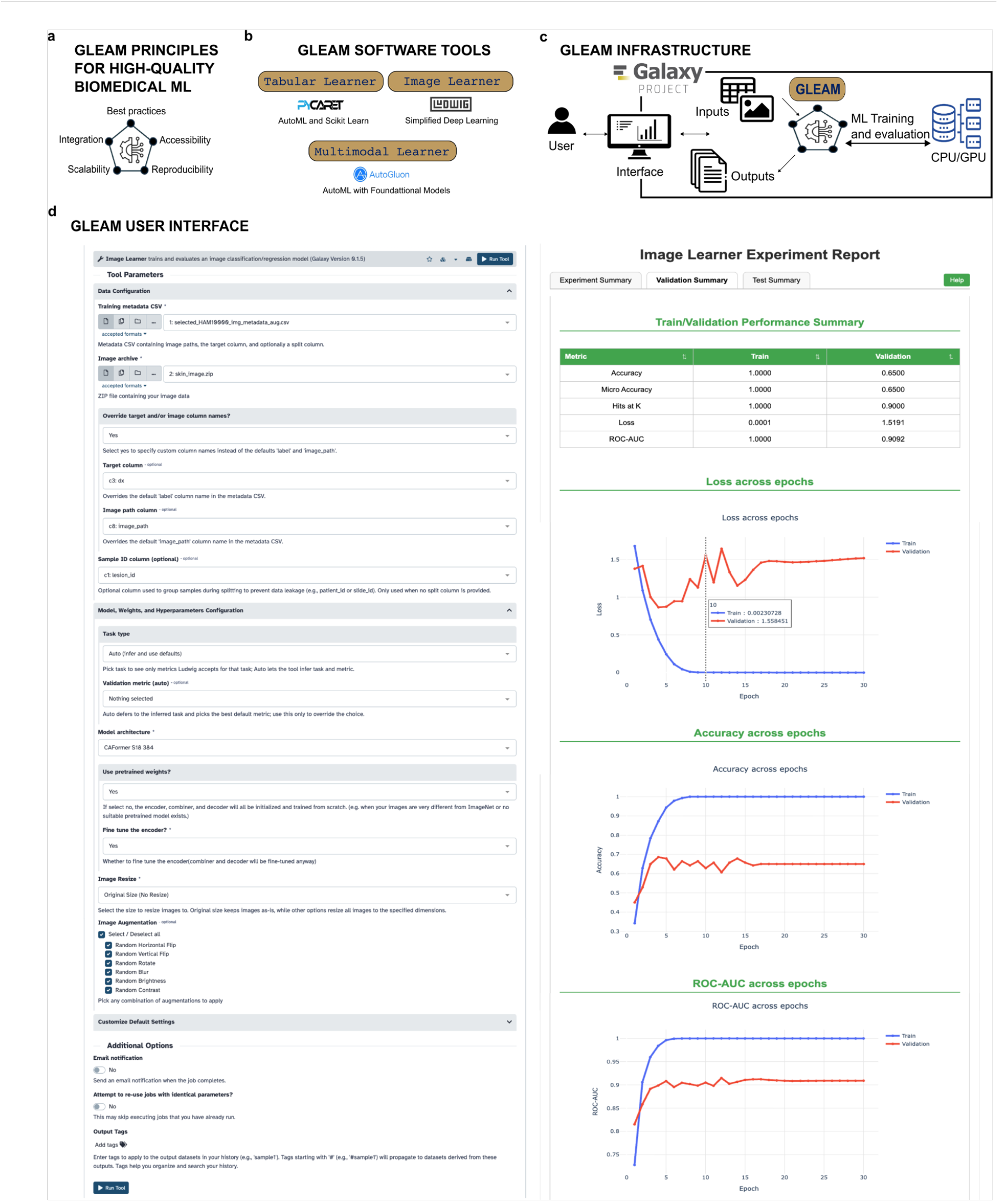
GLEAM principles, toolkit composition, and Galaxy-based execution workflow. (a) Five principles informed GLEAM’s implementation: accessibility, integration, reproducibility, scalability, and best practices. (b) The GLEAM toolkit has three learner tools that each leverages a different machine learning software library: `Tabular Learner` uses PyCaret for traditional supervised learning on tabular data, `Image Learner` uses Ludwig for deep learning on images, and `Multimodal Learner` uses AutoGluon for stacked ensemble learning across modalities. (c) Galaxy infrastructure and job flow. Scientists interact with GLEAM through the Galaxy web interface by providing inputs and selecting options, and GLEAM executes training, validation, and held-out test jobs on available CPU or GPU resources and returns outputs to the Galaxy history. (d) Example user workflow showing the web-based learner interface used to supply inputs and configure a run, and the standardized interactive HTML report that summarizes performance metrics and diagnostics and provides per-sample predictions, exported models, and recorded run settings for review and reproducibility.

GLEAM is implemented in Galaxy^21–23^ (https://galaxyproject.org), a popular web-based computational workbench for biomedical research (**Fig. 1c**). Thousands of scientists use Galaxy daily for all kinds of biomedical data analyses, from genomics to image analysis to chemical informatics. We previously developed a suite of Galaxy tools for machine learning^24^, and GLEAM is the next iteration of machine learning tools in Galaxy. With Galaxy and GLEAM, scientists can run modern ML experiments using a simple and transparent user interface regardless of their informatics expertise. All learners have a similar user interface with the same layout and parameter organization (**Fig. 1d; Supplementary Table 4**). Importantly, this interface helps scientists configure their ML analysis. When the analysis is complete, the analysis results and artifacts are viewable in Galaxy’s interface via a web-based report.

Integration with Galaxy provides GLEAM with several benefits. GLEAM learners can be connected to more than 10,000 Galaxy tools spanning genomics, transcriptomics, proteomics, metabolomics, microbiome analysis, imaging, single-cell omics, and related data types, all through a web browser. To support learning and training in biomedical research, the Galaxy Training Network^25^ (https://training.galaxyproject.org/) provides more than 520 hands-on tutorials covering both individual tools and end-to-end workflows. We have developed tutorials for the Galaxy Training Network for GLEAM’s learners. Galaxy runs GLEAM ML analyses on high-performance computing resources when available. For GPU-capable tools, Galaxy can schedule execution on GPUs, so users can scale deep-learning runs without manual cluster setup. Galaxy also standardizes the software environment by managing tool dependencies and versions through Conda packages and containers, reducing installation and version-mismatch errors that commonly undermine reproducibility. GLEAM keeps the run context together with the outputs and supports sharing via link, so collaborators can inspect what was run and repeat analyses under the same settings.

### Predicting Patient Response to Immunotherapy Using Tabular Machine Learning

GLEAM’s `Tabular Learner` tool uses the PyCaret^16^ software library for automated machine learning using tabular datasets. `Tabular Learner` creates and evaluates >15 ML model families, including linear models, kernel methods, tree ensembles, and gradient-boosted methods **(Supplementary Table 5)**. Models are trained with cross-validation and ranked by one of six metrics, such as accuracy, F1 score, or Area Under the Receiver Operating Characteristic Curve (ROC AUC). For binary tasks, users can set the decision threshold on predicted probabilities. To improve transparency, `Tabular Learner` includes SHAP feature importance analyses that quantify how each feature contributes to the predicted outcome across samples **(Supplementary Fig. 1)**. `Tabular Learner` produces the best model and an HTML report with performance metrics, hyperparameters, and feature importance, all recorded in the Galaxy history. The report organizes results into navigable tabs, including Experiment Summary, Validation Summary, Test Summary, and Feature Importance, so users can easily locate key outputs **(Supplementary Fig. 2)**. An interactive, step-by-step tutorial for `Tabular Learner` is available via the Galaxy Training Network^25,26^.

We applied `Tabular Learner` to the Chowell pan-cancer immunotherapy dataset^27^ that was previously analyzed using machine learning to predict patient response to therapy^28^ (**Fig. 2a**). This dataset includes six clinical and genomic features for each patient and a target binary variable indicating whether a patient responded to immunotherapy. We used the `Tabular Learner` with default parameters and this dataset as input, and a logistic regression model most accurately predicted patient response to immunotherapy **(Supplementary Figs. 2 and 3)**. This is consistent with published results. Furthermore, `Tabular Learner` automatically selects the operating point— the model probability threshold separating positive and negative predictions—that optimizes model performance for a specified metric. Operating point is selected using cross-validated training predictions, and the F1 score is the default metric used for optimization. The `Tabular Learner` report summarizes how precision, recall, and F1 score vary across model probability thresholds, making the tradeoff between these metrics explicit before held-out test evaluation (**Fig. 2b**).

**Figure 2.**
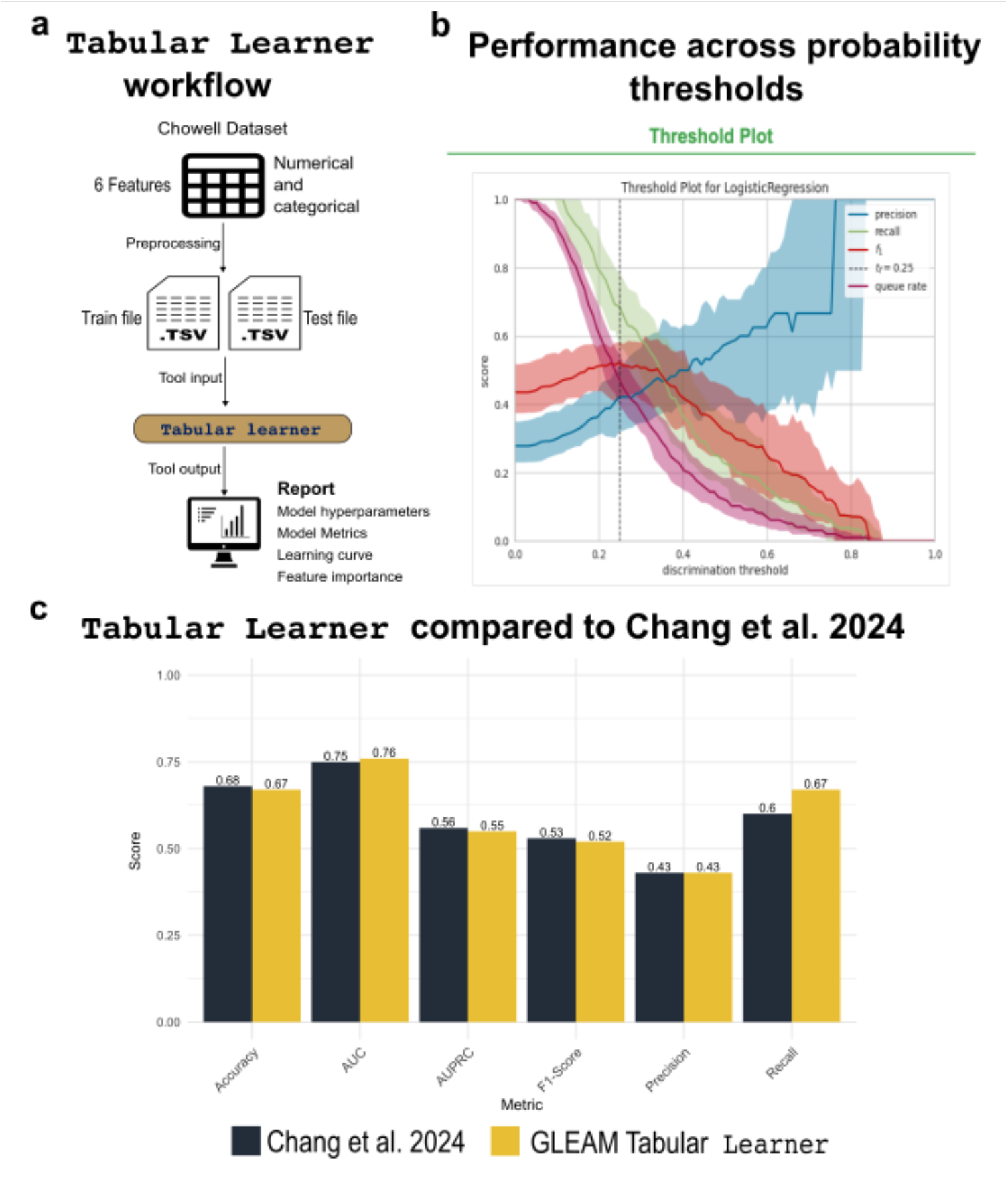
Tabular Learner creates immunotherapy-response model with report-guided operating-point selection. (a) End-to-end `Tabular Learner` workflow in Galaxy using the predefined training and held-out test tables from the LORIS study, from dataset input through standardized report generation. (b) Threshold analysis from the `Tabular Learner` report showing how precision, recall, and F1 score vary across probability cutoffs on the validation set, used to select the decision threshold applied to the test set. (c) Metric-level comparison of the published LORIS benchmark and the `Tabular Learner` model evaluated on the same split and feature set, illustrating comparable discrimination and threshold-dependent performance.

For the Chowell dataset, automatic threshold selection identified that an operating point of 0.269 optimized the F1 score. Using this threshold, `Tabular Learner` produced more balanced held-out test performance than from a 0.5 operating point, with higher recall at the cost of slightly lower precision. On the held-out test set, `Tabular Learner` achieved a ROC AUC of 0.76, an AUPRC of 0.55, a precision of 0.43, a recall of 0.67, and an F1 score of 0.52. These results are nearly identical to the published model, which reported a ROC AUC of 0.75 and an AUPRC of 0.56, and `Tabular Learner` outperformed the published model in recall performance (**Fig. 2c**).

The full analysis history for both executions of the `Tabular Learner`, with evaluation reports, is available at https://usegalaxy.org/u/lyra-jr/h/tabular-learner-immunotherapy-response-prediction (see Methods section for complete history details). These results show that `Tabular Learner` can produce a high-quality ML model while substantially reducing the manual effort typically required to train, document, and evaluate models.

### Skin Lesion Classification With Image-based Machine Learning

GLEAM’s `Image Learner` enables scientists to easily create sophisticated predictive models that use images as input along with a metadata table of image labels and other information. Model types available in `Image Learner` include the 74 standard TorchVision models as well as the 55 MetaFormer foundation image models^29^ **(Supplementary Table 4)**. These models can either be trained from scratch using only the input dataset or be initialized with pretrained weights and then fine-tuned on the input dataset. `Image Learner` supports data augmentation, including random flips, rotations, blurs, and changes to brightness and contrast adjustments to improve model performance. `Image Learner` also includes functionality to prevent data leakage and split data into representative training and test datasets. An explicit split column can be used to divide images between training and test datasets, or a Sample ID column can be provided so that related images are assigned to the same split set. `Image Learner` is implemented using the Ludwig^30^ software library together with custom code. An interactive, guided tutorial on `Image Learner` is available through the Galaxy Training Network^31^.

We applied `Image Learner` to the HAM10000 dermatoscopic image dataset ^32^ to build an ML model that can predict skin lesion diagnosis. The HAM10000 dataset contains 10,015 skin lesion images assigned to seven diagnostic categories, including benign lesions and multiple types of skin cancer^32^. To evaluate `Image Learner` under both benchmarking and real-world conditions, we prepared two versions of the dataset using the same standardized workflow structure composed of a metadata table and an image archive as tool inputs (**Fig. 3a**). First, we reproduced a published class-balanced HAM10000 subset^33^ by selecting 100 images per category and generating horizontally flipped copies, yielding 1,400 images total. Second, we used the complete HAM10000 dataset in its original form. Using the pretrained CAFormer S18 384 foundation model within `Image Learner`, we generated a comprehensive performance report from both datasets that included overall performance, class-wise error patterns, a confusion matrix, and a per-class metrics heatmap (**Supplementary Fig. 4-7**).

In the class-balanced dataset, the confusion matrix and per-class metrics heatmap revealed residual error patterns across the seven diagnostic categories in the held-out test set (**Fig. 3b**). The model achieved a test accuracy of 0.87 and an F1 score of 0.88, performing comparably to published convolutional neural network benchmarks while showing slightly higher recall and F1 scores^33^ (**Fig. 3c**). In the complete HAM10000 dataset, `Image Learner` maintained strong class-level performance, with F1 scores of 0.91 for vascular lesions, 0.95 for melanocytic nevi, and 0.74 for melanoma. These patterns closely matched those reported for a published EfficientNet-B1 architecture^34^ (**Fig. 3d**).

Data leakage is a common risk with biomedical datasets that can lead to incorrectly inflated performance results. Data leakage and inflated performance occur when images from the same sample or cohort appear in both the train and test datasets, allowing the model to use features memorized from the training dataset to artificially inflate performance in the held-out test dataset. In HAM10000, two potential leakage sources are present: (1) paired images created through horizontal flipping and (2) multiple images originating from the same lesion. To assess their impact, we repeated the analysis using the Sample ID column option in the tool to enforce lesion-level partitioning. Leakage-aware partitioning reduced test performance in both HAM10000 analyses, with the larger effect in the complete dataset (**Fig. 3e;** class-balanced subset in **Supplementary Fig. 8**). The largest class-level F1 reductions occurred for actinic keratosis and melanoma, which decreased from 0.69 to 0.46 and from 0.74 to 0.58, respectively (**Fig. 3e**).

**Figure 3.**
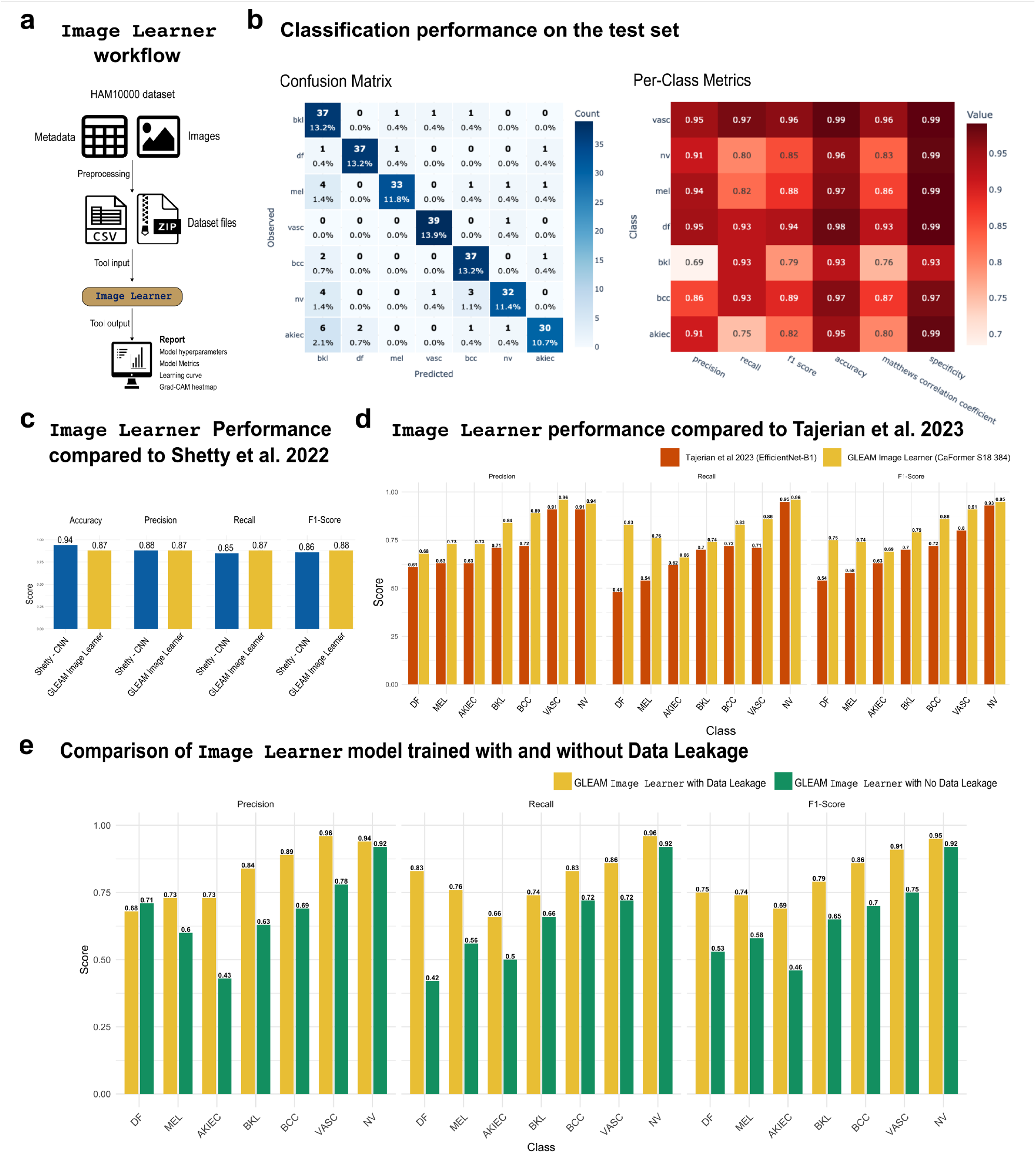
Image Learner benchmarks HAM10000 classification and reveals performance inflation from lesion-level data leakage. (a) End-to-end `Image Learner` workflow in Galaxy. A sample-level metadata table links target labels to image filenames, and a single ZIP archive provides the images. The tool trains a model to predict labels using images and produces a standardized report; (b) Confusion matrix and per-class metric heatmap on the held-out test set from HAM10000, summarizing predictions across the seven HAM10000 diagnostic classes: dermatofibroma (df), melanoma (mel), actinic keratoses or intraepithelial carcinoma (akiec), benign keratosis-like lesions (bkl), basal cell carcinoma (bcc), vascular lesions (vasc), and melanocytic nevi (nv); (c) Overall performance comparison between `Image Learner` using a CAFormer backbone and the CNN baseline reported by Shetty et al. 2022, evaluated under the same class-balanced subset protocol with horizontal flipping; (d) Per-class precision, recall, and F1 score for `Image Learner` compared with the EfficientNet-B1 benchmark reported by Tajerian et al. 2023, showing similar class-level patterns across categories. (e) Impact of leakage-aware splitting on the full HAM10000 dataset. Metrics from `the Image Learner` run shown in panel d are compared with an otherwise matched run using lesion-aware partitioning, which assigns related images from the same lesion to the same dataset partition. Performance decreased across diagnostic classes, indicating that naive image-level splits can inflate test-set estimates on HAM10000.

The complete analysis histories for the two `Image Learner` models, including the full evaluation reports, are available at https://usegalaxy.org/u/lyra-jr/h/image-learner-ham10000-lesion-classification. These results show that identifier-aware splitting is an important best practice for fair evaluation when there are multiple related images. By providing data splitting functionality, GLEAM makes leakage-aware partitioning a routine part of image-model development, ensuring that reported performance accounts for and mitigates potential data-leakage-induced performance inflation.

### Using Multimodal Machine Learning to Predict Head and Neck Cancer Recurrence

GLEAM’s `Multimodal Learner` trains and evaluates predictive deep learning models from a combination of structured and unstructured inputs. Users provide a single sample-level table whose feature columns can include numeric, categorical, image, and free-text values. Images are provided to the `Multimodal Learner` as a ZIP file. Using the `Multimodal Learner`, users can choose from eight text embedding models from Hugging Face Transformers^35^ and 76 image backbone models from the PyTorch Image Models (TIMM)^36^ library **(Supplementary Table 5)**. `Multimodal Learner` applies the selected encoders to corresponding feature types and combines feature embeddings using a multilayer perceptron fusion head to learn cross-modal interactions and output predictions. Because this end-to-end multimodal training is computationally intensive and can be sensitive to nondeterminism in GPU execution, `Multimodal Learner` provides an optional deterministic mode to support reproducible runs and best-practice reporting. This mode forces PyTorch and CUDA to use deterministic algorithms and fixed seeds, making runs more reproducible at the cost of speed. Model performance is reported using an HTML report that records the full experiment context, including parameter settings, selected hyperparameters, and plotted figures, and makes it available through the Galaxy history. `Multimodal Learner` is implemented using the AutoGluon^37^ software library and custom code for its final report. The Galaxy Training Network provides a tutorial on `Multimodal Learner` to better understand the tool setup and usage^38^.

We applied `Multimodal Learner` to predict cancer recurrence in the HANCOCK dataset^39^, a cohort of 763 head and neck cancer patients that includes structured clinical variables, unstructured clinical text, and histologic images. For multimodal integration, we constructed a single input table comprising clinical variables, a text column containing International Classification of Diseases (ICD) codes, and two image-path columns corresponding to CD3- and CD8-stained tissue microarray images. A separate image archive containing all CD3 and CD8 images was linked to individual patients through these paths (**Fig. 4a**). This input format contrasts with the original HANCOCK modeling pipeline, which used engineered features derived from the same modalities. In the original workflow, histologic images were processed to quantify CD3 and CD8 cell positivity, and clinical free text was converted into a bag-of-words representation, reducing all inputs to structured numerical variables before model training. By retaining text and images in their original form, `Multimodal Learner` reduces the need for manual feature engineering while preserving the predefined training and held-out test split from the published study. In this configuration, the `Multimodal Learner` text and image embedding models encode each modality into its own embedding, and late fusion is used to integrate all patient data to produce a patient recurrence prediction score (**Fig. 4b**).

**Figure 4.**
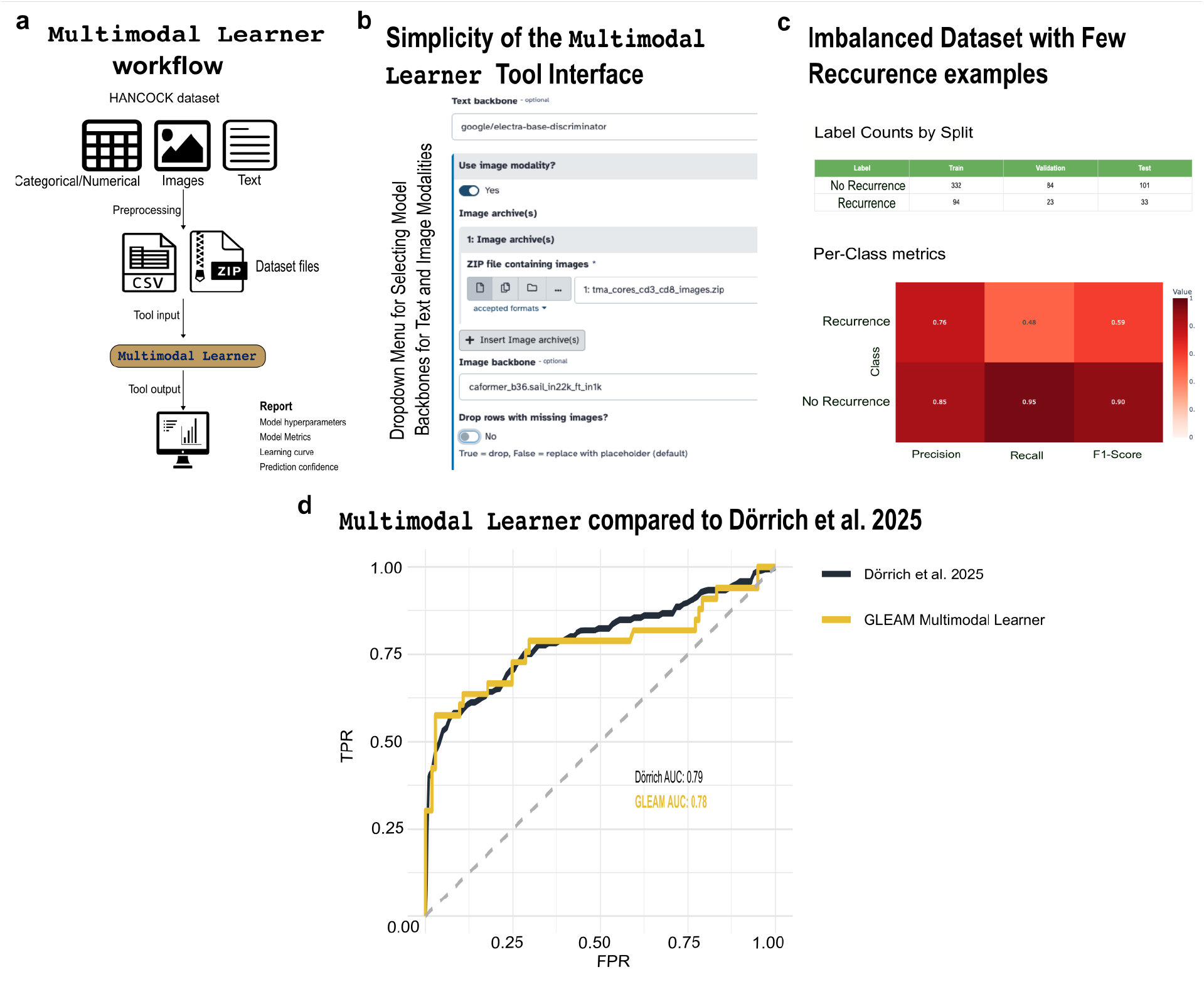
Multimodal Learner predicts head and neck cancer recurrence from clinical variables, ICD text, and paired histology images. (a) input structure and workflow for `Multimodal Learner`. A single sample-level table links each patient to structured clinical fields, an ICD text column, and separate CD3 and CD8 image columns. Images are provided in a single ZIP archive. `Multimodal Learner` produces a standardized report and machine-readable outputs. (b) `Multimodal Learner` user interface, showing that an end-to-end multimodal experiment can be configured by selecting the input table, modality columns, target label, and training preset without writing code. (c) Report summaries on the held-out test set, including label counts across train, validation, and test splits and class-specific precision, recall, and F1 scores for recurrence and no recurrence. (d) ROC curve comparison between `Multimodal Learner` results and the published HANCOCK benchmark on the same predefined split, showing closely matched performance. True Positive Rate (TPR); False Positive Rate (FPR).

The HANCOCK dataset is imbalanced, with far fewer recurrence cases than no-recurrence cases, so we evaluated performance of the `Multimodal Learner` at the class level in addition to aggregate metrics (**Fig. 4c**; **Supplementary Fig. 9 and 10**). On the held-out test set, the `Multimodal Learner` model achieved a precision of 0.85, a recall of 0.95, and an F1 of 0.90 for the *no recurrence* class, versus 0.76, 0.48, and 0.59 for the *recurrence* class (**Fig. 4c**). The published HANCOCK baseline reported a ROC AUC of 0.79^39^, whereas `Multimodal Learner` achieved a comparable ROC AUC of 0.78 with similar ROC curve behavior across the operating range (**Fig. 4d**). These results indicate that the `Multimodal Learner` performance is comparable to prior approaches while operating directly on multimodal inputs, without additional feature engineering. The complete analysis history, including the input datasets and output files, is available at https://usegalaxy.org/u/lyra-jr/h/multimodal-learner-head-and-neck-recurrence-predictor. Together, these results show that `Multimodal Learner` enables rapid development of high-quality multimodal predictive models using a code-free framework and provides insight into class-specific strengths and weaknesses through its evaluation report.

## DISCUSSION

Today’s software tools do not make it easy for biomedical researchers to use machine learning to its fullest capabilities. Our Galaxy Learning and Modeling (GLEAM) software toolkit fills this key gap by empowering all scientists to perform complete and robust machine learning analyses regardless of their informatics expertise. GLEAM’s three learners—`Tabular Learner`, `Image Learner`, and `Multimodal Learner`—enable end-to-end machine learning for the most common datatypes in biomedicine without the need to write code. GLEAM’s integration with the Galaxy computational workbench is essential as Galaxy provides key functionality for GLEAM. GLEAM uses Galaxy to provide a guided web interface for data inputs and parameter settings as well as a comprehensive report for evaluating ML experiments. GLEAM and Galaxy use AutoML software libraries and CPU or GPU resources to create state-of-the-art ML models. GLEAM reports are web-based and summarize data inputs, trained models, predictions, and model results in one place. In several analyses run with GLEAM’s learners, we demonstrate how GLEAM can advance use of machine learning in biomedicine. This includes explicit threshold selection in the `Tabular Learner` to balance multiple metrics, evaluating and mitigating data leakage in the `Image Learner`, and using foundation models on multimodal datasets in the `Multimodal Learner`. Across the three use cases, GLEAM produced results comparable to published benchmarks while greatly simplifying entire analyses and preserving analysis traceability through Galaxy histories.

GLEAM addresses critical challenges in biomedical machine learning that others have documented. Median adherence to the Transparent Reporting of a multivariable prediction model for Individual Prognosis or Diagnosis (TRIPOD) reporting items in biomedical machine learning prediction studies was 38.7%, and key elements were often missing, including final model specification, instructions for applying the model, and a complete description of predictive performance^40,41^. The Prediction Model Risk of Bias Assessment Tool (PROBAST)+AI recommends reporting separate model development from model performance evaluation^42^. Data leakage remains a major driver of overoptimistic claims when preprocessing crosses the boundary between training and test data or when data splits do not respect dependence between samples^43^. GLEAM helps scientists address these issues by enforcing intentional data partitioning, documenting all inputs and models used in a machine learning analysis, and generating standardized reports that capture configuration, software versions, and class-specific performance summaries.

GLEAM fills a unique space in the software ecosystem that enables biomedical machine learning. Computational workbenches such as Galaxy, GenePattern^44^, Terra (https://app.terra.bio/), and the NCI Cancer Cloud Resources^45^ democratize biomedical analyses and enable anyone to perform analyses. These workbenches integrate a graphical user interface, workflow engine, dataset management, and computing resources to provide a single place for scientists to perform data analyses. GLEAM extends Galaxy to enable comprehensive and best practice machine learning. Importantly, GLEAM’s tools can be used with the tens of thousands of tools that have already been integrated with Galaxy^46^ to integrate machine learning into other biomedical data analyses.

Despite these strengths, GLEAM has several limitations. GLEAM enables common machine learning analyses but does not replace custom development. New methods, neural architectures, and training objectives cannot be accomplished with GLEAM. GLEAM is also dependent on the underlying AutoML software libraries, which restricts the ML models and parameter settings that can be used. While GLEAM steers users towards the use of best practices in machine learning, scientists can still use GLEAM in ways that violate best practices. Appropriate data collection and splitting, external validation, dataset shift assessment, fairness audits, and other best practices in machine learning remain the responsibility of the scientific community.

Future work will expand GLEAM in several directions. We will add learners for additional data modalities and incorporate foundation models into existing and new learners. We will also incorporate automated checks to help scientists catch common threats to ML model validity, including leakage, incomplete dataset documentation, and calibration shifts. Finally, we will extend GLEAM’s evaluation reports with additional statistical rigor such as confidence intervals and calibration metrics. These directions adhere to GLEAM’s core goals and leverage evolving technological and statistical advances to improve biomedical machine learning.

## ONLINE METHODS

All analyses were executed in Galaxy, which records input datasets, tool versions, and parameter settings in the history. Each GLEAM run returns an interactive report, per-sample predictions, saved model artifacts, and a structured run configuration. Together, these outputs and the Galaxy history provide a complete record that supports review and reruns with the same inputs and settings.

### Tabular Learner: LORIS dataset acquisition, preprocessing, and model configuration

`Tabular Learner` performs supervised learning for structured datasets with automated preprocessing, model training, and standardized reporting. Preprocessing includes missing-value handling, categorical encoding, and optional normalization. Models are compared under stratified cross-validation, with optional hyperparameter optimization and probability calibration as configured by the user. Outputs include an interactive HTML report with summary metrics, ROC and precision– recall curves, confusion matrices, calibration and threshold analyses, together with saved model artifacts, run settings, and per-sample predictions.

For the immunotherapy response analysis, we used the publicly released AllData.xlsx file, distributed with the LORIS study, which was derived from the multi-cancer immune checkpoint blockade cohort compiled by Chowell and colleagues. We extracted the predefined training and held-out test partitions, retaining the predictors used in the published LORIS model: tumor mutational burden, age, neutrophil-to-lymphocyte ratio, serum albumin, systemic therapy history, and one-hot indicators for cancer type. The binary label Response was defined as an objective response, equal to 1, and a non-response, equal to 0. To match the published preprocessing, extreme values were clipped to the reported thresholds, with tumor mutational burden capped at 50, age capped at 85, and neutrophil-to-lymphocyte ratio capped at 25. Processed training and test partitions were exported as tab-separated files and provided as direct inputs to `Tabular Learner`.

`Tabular Learner` was run in classification mode using the formatted training table for model development and the held-out test table for final evaluation, with the Response column specified as the target for prediction. Model selection used the tool’s default cross-validation and search settings, as recorded in the Galaxy history and configuration output. For binary classification, `Tabular Learner` evaluated candidate operating points—potential model probability thresholds separating positive and negative predictions—using cross-validated training predictions and selected the threshold that optimized the user-specified metric. The default metric, F1 score, was used to select the optimal operating point. `Tabular Learner` then applied the selected operating point, 0.269, to the make predictions on the held-out test set. The threshold marker shown in the report plot may differ slightly from the selected decision threshold because the visualization and optimizer may use different threshold grids, rounding rules, or tie-breaking rules for near-equivalent metric values.

### Image Learner: HAM10000 dataset acquisition, preprocessing, and configuration

For dermoscopic image classification, we utilized the HAM10000 dataset, which comprises 10,015 dermoscopic images categorized into seven diagnostic categories: akiec, bcc, bkl, df, mel, nv, and vasc. Images and metadata were obtained from the public distribution and filtered to retain samples with a single unambiguous class label.

To align with the class-balanced subset protocol used in the published benchmark, we selected 100 images per category and created one horizontally flipped counterpart for each image, yielding 200 images per class and a total of 1,400 images. We generated a sample-level table with two required fields: a categorical target column named label and an image path column that matches filenames in the image archive. The corresponding images were provided as a single ZIP archive. No predefined split column was supplied, so `Image Learner` created stratified partitions using tool defaults, which are recorded in the run configuration.

Within Galaxy, `Image Learner` was configured for multiclass classification with label as the target. The pretrained model was CAFormer-S18 with 384-pixel input resolution. Training was run for 30 epochs with a batch size of 32. Images were provided in their stored resolution and resized internally to the expected input size during training. Unless otherwise specified, augmentation and early stopping used tool defaults, and all settings were recorded in the Galaxy history and report outputs.

Because HAM10000 includes multiple images from the same lesion in parts of the collection, and external augmentation can create near-duplicate views, we also performed a leakage-aware sensitivity run using two `Image Learner` options. We provided an identifier column to group related samples so that automated splitting keeps all images from the same lesion within a single partition. We also enabled augmentation within `Image Learner`, so augmented variants are generated during training rather than introduced as additional files before splitting. Results from this leakage-aware run are present in the Galaxy history.

### Multimodal Learner: HANCOCK dataset acquisition and processing

`Multimodal Learner` trains late-fusion prediction models from heterogeneous biomedical inputs using AutoGluon Multimodal. Each experiment is defined by a sample-level table that includes the outcome label and columns for one or more modalities from the same individual, including tabular variables and, optionally, text and images. The tool trains modality-specific encoders and combines their representations to produce a final predictor. Outputs include an interactive report with performance summaries and diagnostic plots, per-sample predictions, and a structured run configuration.

For the analysis of head and neck cancer recurrence, we utilized the HANCOCK resource and the predefined train and test splits from the published study. We assembled separate training and testing tables that included the binary recurrence label as the target, clinical covariates as both categorical and numerical fields, an ICD text column retained as plain text, and two image filename columns that link each sample to CD3 and CD8 tissue microarray core images. All CD3 and CD8 images were provided as a single ZIP archive that matches the filenames referenced in the tables. The train and test tables were provided to `Multimodal Learner` as separate development and held-out evaluation inputs. An internal validation split was created from the training table using tool defaults.

`Multimodal Learner` was configured with ELECTRA-base as the text embedding model and CAFormer-B36 as the image embedding model. The ROC AUC was used as the primary metric for model selection, and the random seed was set to 42. Missing-image row dropping was disabled. A binary decision threshold of 0.29 was specified for threshold-dependent metrics. All embedding models, optimization settings, and evaluation parameters were captured in the configuration output and the Galaxy history.

## Code and data availability

A website summarizing GLEAM, with links to its tools, tutorials, and source code, is available at https://goeckslab.github.io/gleam/.

Code repositories for GLEAM tools are hosted on GitHub, and tool versioning and distribution are managed through the Galaxy ToolShed^46^. The source code and preprocessing resources for the use cases are available at https://github.com/goeckslab/gleam and https://github.com/goeckslab/gleam_use_cases.

Complete Galaxy histories for all analyses, including inputs, parameters, and outputs, are publicly available. `Tabular learner` history: https://usegalaxy.org/u/lyra-jr/h/tabular-learner-immunotherapy-response-prediction, `Image learner` history: https://usegalaxy.org/u/lyra-jr/h/image-learner-ham10000-lesion-classification, and `Multimodal learner` history: https://usegalaxy.org/u/lyra-jr/h/multimodal-learner-head-and-neck-recurrence-predictor.

Datasets used in each experiment are available on Zenodo. For the `Tabular Learner` immunotherapy response task: https://zenodo.org/records/17781688. For the `Image Learner` skin lesion classification task: https://zenodo.org/records/18284218. For the `Multimodal Learner` head and neck cancer recurrence task: https://zenodo.org/records/18603388.

## Software and Hardware

All GLEAM analyses were run in Galaxy using containerized environments defined in the Galaxy tool XML wrappers. The `Tabular Learner` uses Galaxy tool profile 21.05, wrapper version 0.1.5, and quay.io/goeckslab/galaxy-pycaret:3.3.2, which provides PyCaret 3.3.2 and its dependencies. The `Image Learner` and `Multimodal Learner` use Galaxy tool profile 22.01. The `Image Learner` uses wrapper version 0.1.5 and quay.io/goeckslab/galaxy-ludwig-gpu:0.10.1, which provides Ludwig 0.10.1 with GPU-enabled deep learning dependencies. The `Multimodal Learner` uses wrapper version 0.1.9 and quay.io/goeckslab/multimodal-learner:1.4.0, which provides the AutoGluon multimodal stack and supporting libraries. Galaxy executes these containers through the deployment runtime, such as Docker, Singularity, or Apptainer.

## Use of large language models

OpenAI Codex 5.5 was used to assist with writing and refining analysis code. All generated outputs were reviewed and verified by the authors, who take full responsibility for the final content.

