## Supplemental figures for "A Web-based software toolkit for accessible and best-practice machine learning analyses in biomedical research"

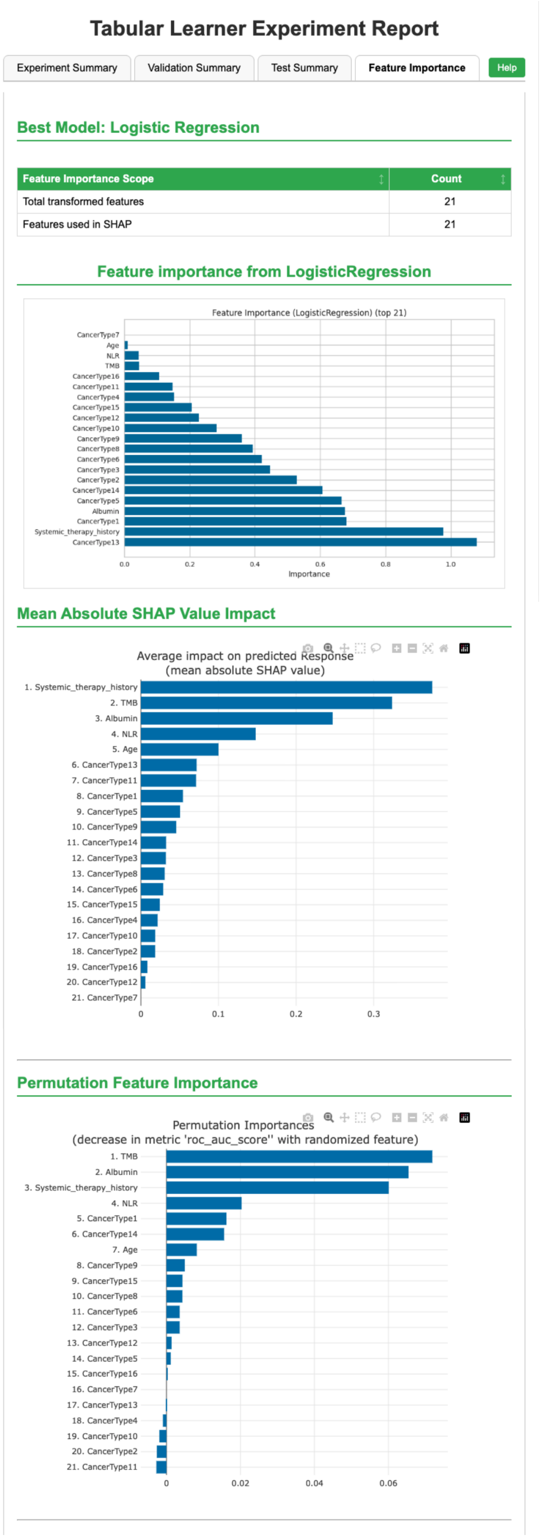


**Supplementary Fig. 1 | Feature importance for Tabular Learner prediction of immunotherapy response.** Feature-importance output for the logistic regression model selected by Tabular Learner in the immunotherapy response task. Variables are ranked using model-based importance, mean absolute SHAP value and permutation importance, providing complementary views of the features that contributed most to prediction.

### Tabular Learner Experiment Report

Experiment Summary | Validation Summary | Test Summary | Feature Importance | Help

#### Best Model: Logistic Regression

#### Dataset Overview

| Label | Training | Test |
|---|---|---|
| 0 | 682 | 388 |
| 1 | 282 | 127 |

#### Best Model Performance

| Metric | Train | Validation | Test |
|---|---|---|---|
| Accuracy | 0.663 | 0.655 | 0.697 |
| ROC-AUC | 0.746 | 0.710 | 0.760 |
| Precision | 0.450 | 0.441 | 0.427 |
| Recall | 0.691 | 0.677 | 0.669 |
| F1-Score | 0.545 | 0.534 | 0.521 |
| PR-AUC | 0.551 | 0.515 | 0.553 |
| Specificity | 0.651 | 0.645 | 0.706 |
| MCC | 0.313 | 0.295 | 0.332 |

#### Experiment and Data Parameters

| Parameter | Value |
|---|---|
| Best Model Metric | Accuracy |
| Target | Response |
| Random Seed | 42 |
| Train Size | 964 rows |
| Normalize | False |
| Feature Selection | False |
| Cross Validation | True |
| Cross Validation Folds | 10 |
| Remove Outliers | False |
| Remove Multicollinearity | False |
| Polynomial Features | False |
| Models | None |
| Fix Imbalance | False |
| Threshold source | Optimized on cross-validated training predictions |
| Threshold optimization metric | F1 |
| Decision threshold (Test) | 0.269 |

#### Best Model Hyperparameters

| Parameter | Value |
|---|---|
| C | 1.0 |
| Class Weight | None |
| Dual | False |
| Fit Intercept | True |
| Intercept Scaling | 1 |
| L1 Ratio | None |
| Max Iter | 1000 |
| Multi Class | auto |
| N Jobs | None |
| Penalty | l2 |
| Random State | 42 |
| Solver | lbfgs |
| Tol | 0.0001 |
| Verbose | 0 |
| Warm Start | False |

**Supplementary Fig. 2 | Tabular Learner configuration for automatic decision-threshold selection.** Configuration report for the immunotherapy response analysis using automatic threshold selection. The report records the input dataset, target label, selected logistic regression model, evaluation setup, threshold-optimization metric and selected decision threshold. Tabular Learner optimized F1 score using cross-validated training predictions and selected a probability threshold of 0.269, which was then applied to the held-out test set.

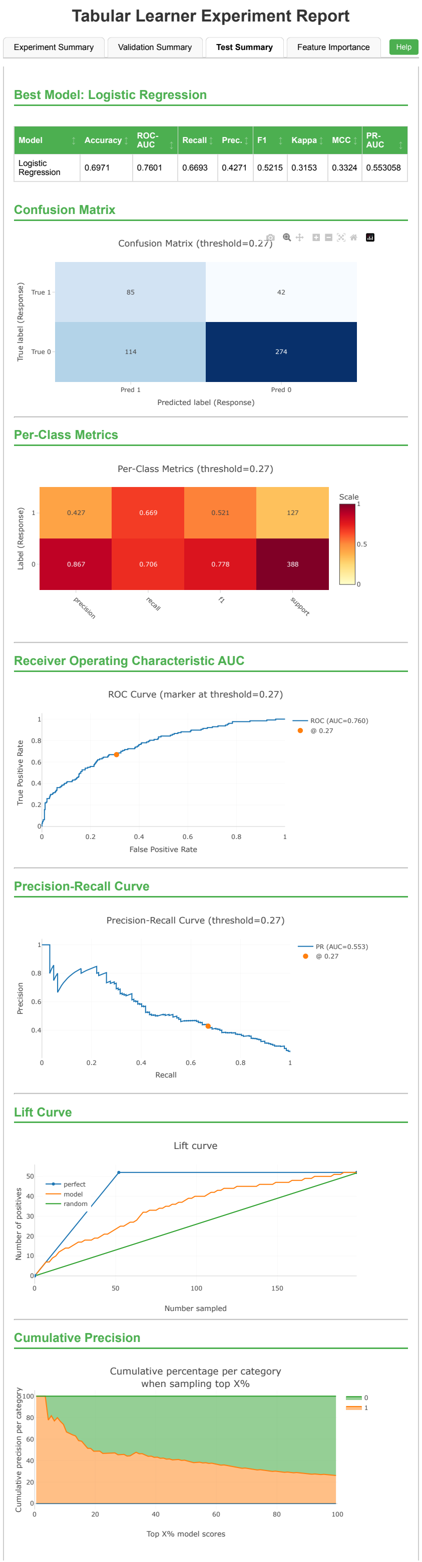


**Supplementary Fig. 3 | Test performance of Tabular Learner using the automatically selected decision threshold.** Test Summary report for the selected logistic regression model evaluated on the held-out test set after automatic threshold selection. Tabular Learner optimized F1 score using cross-validated training predictions and selected a probability threshold of 0.269. At this operating point, the model achieved a ROC AUC of 0.760, AUPRC of 0.553, precision of 0.427, recall of 0.669 and F1 score of 0.521 on the held-out test set.

#### Image Learner Experiment Report

Experiment Summary | Validation Summary | Test Summary | Help

##### Dataset Overview

| Label | Train | Validation | Test |
|---|---|---|---|
| akiec | 140 | 20 | 40 |
| bcc | 140 | 20 | 40 |
| bkl | 140 | 20 | 40 |
| df | 140 | 20 | 40 |
| mel | 140 | 20 | 40 |
| nv | 140 | 20 | 40 |
| vasc | 140 | 20 | 40 |

##### Model Performance Summary

| Metric | Train | Validation | Test |
|---|---|---|---|
| Accuracy | 0.9969 | 0.8857 | 0.8750 |
| Micro Accuracy | 0.9969 | 0.8857 | 0.8750 |
| Hits at K | 1.0000 | 0.9786 | 0.9786 |
| Loss | 0.0100 | 1.3686 | 0.5856 |
| ROC-AUC | 1.0000 | 0.9833 | 0.9842 |

##### Training Configuration (Model, Data, Metrics)

| Parameter | Value |
|---|---|
| Architecture | caformer_s18_384 |
| Image Size | 96x96 |
| Target Column | dx |
| Task Type | classification |
| Validation Metric | Accuracy |
| Loss Function | softmax_cross_entropy |
| Epochs | 30 |
| Total Epochs | 30 |
| Batch Size | auto |
| Fine Tune | True |
| Use Pretrained | True |
| Optimizer | adam |
| Random Seed | 42 |
| Early Stop | 30 |
| Use Mixed Precision | No |
| Gradient Clipping | 0.5 |
| Augmentation | random_rotate, random_blur, random_brightness, random_contrast, random_vertical_flip |
| Data Split | No split column in CSV. Created stratified random split: [70, 10, 20]% for train/val/test with balanced label distribution. |

Model trained using Ludwig. Ludwig documentation provides detailed information about default model and training parameters

**Supplementary Fig. 4 | Image Learner configuration for the class-balanced HAM10000 subset without lesion-level grouping.** Configuration report for the class-balanced HAM10000 benchmark subset, which included 1,400 dermatoscopic images after horizontal flipping. The report records class balance, training settings, the pretrained CAFormer S18 384 backbone and the stratified train, validation and test split used without grouping related lesion images.

#### Image Learner Experiment Report

Experiment Summary | Validation Summary | Test Summary | Help

##### Dataset Overview

| Label | Train | Validation | Test |
|---|---|---|---|
| akiec | 229 | 33 | 65 |
| bcc | 360 | 51 | 103 |
| bkl | 769 | 110 | 220 |
| df | 81 | 11 | 23 |
| mel | 779 | 111 | 223 |
| nv | 4694 | 670 | 1341 |
| vasc | 99 | 14 | 29 |

##### Model Performance Summary

| Metric | Train | Validation | Test |
|---|---|---|---|
| Accuracy | 0.9841 | 0.7423 | 0.8044 |
| Micro Accuracy | 0.9919 | 0.8930 | 0.8947 |
| Hits at K | 0.9993 | 0.9323 | 0.9605 |
| Loss | 0.0245 | 0.7539 | 0.5035 |
| ROC-AUC | 0.9999 | 0.9701 | 0.9825 |

##### Training Configuration (Model, Data, Metrics)

| Parameter | Value |
|---|---|
| Architecture | caformer_s18_384 |
| Image Size | 450x600 |
| Target Column | dx |
| Task Type | classification |
| Validation Metric | Accuracy |
| Loss Function | softmax_cross_entropy |
| Epochs | 30 |
| Total Epochs | 30 |
| Batch Size | auto |
| Fine Tune | True |
| Use Pretrained | True |
| Optimizer | adam |
| Random Seed | 42 |
| Early Stop | 30 |
| Use Mixed Precision | No |
| Gradient Clipping | 0.5 |
| Augmentation | random_horizontal_flip, random_vertical_flip, random_rotate, random_blur, random_brightness, random_contrast |
| Data Split | No split column in CSV. Created stratified random split: [70, 10, 20]% for train/val/test with balanced label distribution. |

Model trained using Ludwig. Ludwig documentation provides detailed information about default model and training parameters

**Supplementary Fig. 6 | Image Learner configuration for the full HAM10000 dataset without lesion-level grouping.** Configuration report for the full HAM10000 analysis of 10,015 dermatoscopic images across seven diagnostic classes. The report records dataset composition, training settings, image preprocessing, augmentation settings and fine-tuning of the pretrained CAFormer S18 384 backbone. The train, validation and test split was generated without grouping related lesion images.

#### Image Learner Experiment Report

Experiment Summary | Validation Summary | **Test Summary** | Help

##### Test Performance Summary

| Metric | Test |
|---|---|
| Accuracy | 0.8044 |
| Micro Accuracy | 0.8947 |
| Macro F1-score | 0.8122 |
| Micro F1-score | 0.8947 |
| Weighted F1-score | 0.8937 |
| Macro Precision | 0.8248 |
| Micro Precision | 0.8947 |
| Weighted Precision | 0.8947 |
| Macro Recall | 0.8044 |
| Micro Recall | 0.8947 |
| Weighted Recall | 0.8947 |
| Hits at K | 0.9605 |
| Cohen's Kappa | 0.7956 |
| Loss | 0.5035 |
| ROC-AUC | 0.9825 |
| Token Accuracy | 0.8947 |

##### Confusion Matrix

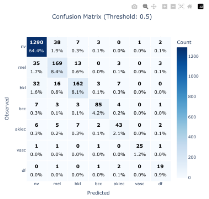


##### Per-Class metrics

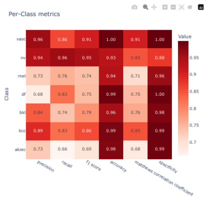


##### Per-Class Metrics (Test)

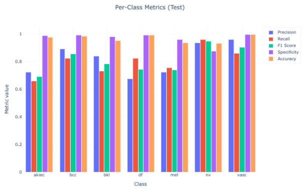


##### Prediction Confidence Distribution

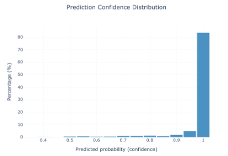


##### Grad-CAM Heatmaps

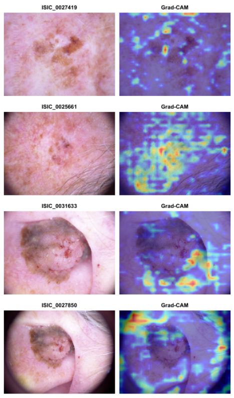


**Supplementary Fig. 7 | Test performance on the full HAM10000 dataset without lesion-level grouping.** Test Summary report for the Image Learner model trained on the full HAM10000 dataset. The report shows held-out test metrics, confusion patterns, per-class performance, confidence distributions and Grad-CAM examples across the seven diagnostic classes. The model reached a held-out test accuracy of 0.80 and micro F1 score of 0.89 under a split that did not group related lesion images.

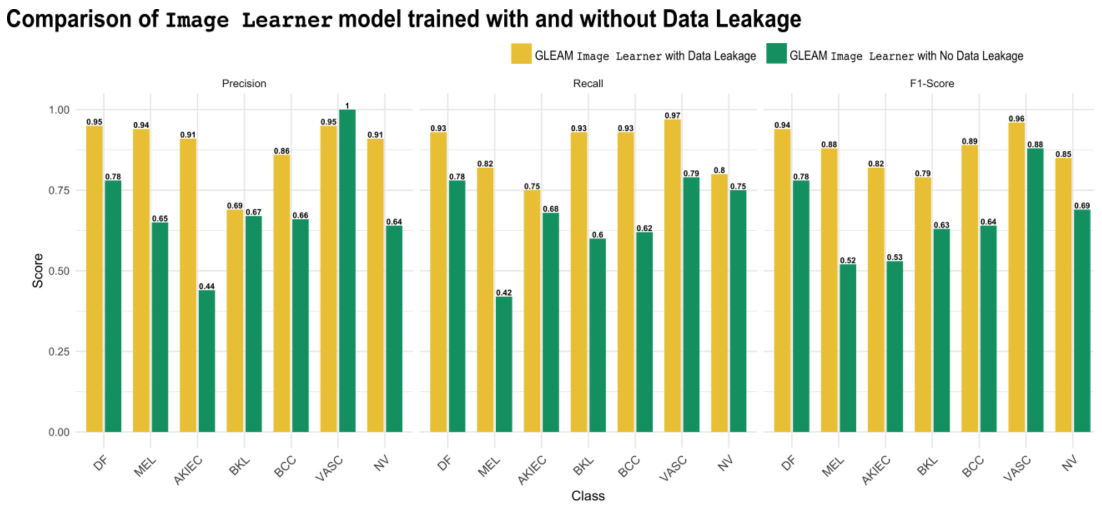


**Supplementary Fig. 8 | Lesion-level grouping reduces apparent HAM10000 class-balanced subset performance.** Held-out test precision, recall and F1 score are shown for each HAM10000 diagnostic class under stratified random splitting and lesion-level grouping. Grouping related images into the same partition reduced performance across diagnostic classes, showing that related lesion images split across training and test partitions can inflate performance estimates.

#### Multimodal Learner Experiment Report

Experiment Summary | Validation Summary | Test Summary | Help

##### Dataset Overview

**Label Counts by Split**

| Label | Train | Validation | Test |
|---|---|---|---|
| 0 | 332 | 84 | 101 |
| 1 | 94 | 23 | 33 |

##### Model Performance Summary

| Metric | Train | Validation | Test |
|---|---|---|---|
| Accuracy | 0.8803 | 0.8598 | 0.8358 |
| ROC-AUC | 0.8390 | 0.7950 | 0.7807 |
| Precision | 0.8909 | 0.7500 | 0.7619 |
| Recall | 0.5213 | 0.5217 | 0.4848 |
| F1-Score | 0.6577 | 0.6154 | 0.5926 |
| PR-AUC | 0.7090 | 0.6415 | 0.6995 |
| Specificity | 0.9819 | 0.9524 | 0.9505 |
| MCC | 0.6223 | 0.5461 | 0.5159 |
| LogLoss | 0.3575 | 0.4009 | 0.4248 |

##### Model Configuration

| Key | Value |
|---|---|
| Target column | target |
| Model evaluation metric | roc_auc |
| Experiment quality | medium_quality |
| Decision threshold (Test) | 0.426 |
| Threshold source | Optimized on validation split |
| Threshold optimization metric | F1 |
| Random seed | 42 |
| Deterministic mode | True |
| GPU count | 1 |
| Training worker count | 8 |
| Evaluation worker count | 8 |
| Model components | ft_transformer, fusion_mlp, hf_text, timm_image |
| Tabular backbone | ft_transformer (categorical, numerical) |
| Image backbone | caformer_b36.sail_in22k_ft_in1k |
| Text backbone | google/electra-base-discriminator |
| Fusion backbone | fusion_mlp |
| Optimizer | adamw |
| Learning rate | 0.0004 |
| Weight decay | 0.001 |
| LR decay | 0.9 |
| Max epochs | 20 |
| Max steps | -1 |
| Early-stop patience | 10 |
| Val check every N epochs | 1 |
| Top K checkpoints | 3 |
| Top K averaging | greedy_soup |
| Global batch size | 128 |

**Supplementary Fig. 9 | Multimodal Learner configuration for recurrence prediction in the HANCOCK cohort.** Configuration report for recurrence prediction from structured clinical variables, ICD text and paired CD3 and CD8 histology images. The report records the tabular, text, image and fusion modules, selected backbones, threshold settings, training parameters and evaluation metrics used for the HANCOCK analysis.

Model Metric Summary and Config | Train and Validation Summary | **Test Summary** | Help

##### Test Performance Summary

| Metric | Test |
|---|---|
| Accuracy | 0.858209 |
| Precision | 0.850000 |
| Recall (Sensitivity/TPR) | 0.515152 |
| F1-Score | 0.641509 |
| Specificity (TNR) | 0.970297 |
| ROC-AUC | 0.784428 |
| PR-AUC | 0.722695 |
| LogLoss | 0.398082 |
| MCC | 0.586940 |

##### Confusion Matrix

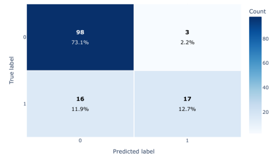


##### Per-Class Metrics

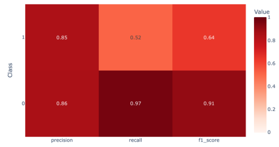


##### ROC Curve

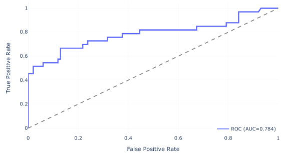


##### Precision–Recall Curve

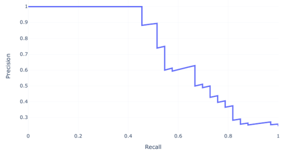


##### Prediction Confidence (Test)

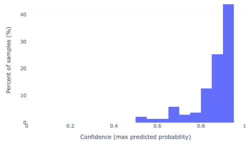


**Supplementary Fig. 10 | Test performance of Multimodal Learner for HANCOCK recurrence prediction.** Test Summary report for the HANCOCK recurrence analysis. The report shows held-out test metrics, confusion patterns between recurrence and no recurrence, class-level precision, recall and F1 score, ROC and precision-recall curves and prediction confidence distributions. On the held-out test set, no recurrence achieved precision, recall and F1 score of 0.85, 0.95 and 0.90, while recurrence achieved 0.76, 0.48 and 0.59. The model achieved a ROC AUC of 0.78 on the predefined test partition.
